# IDEAL-GENOM: Integrated Downstream Analytical Toolkit for Genomic Analysis

**DOI:** 10.1101/2025.08.27.672528

**Authors:** Luis Giraldo González Ricardo, Amabel M.M. Tenghe, Ashwin Ashok Kumar Sreelatha, Manu Sharma

**Affiliations:** Centre for Genetic Epidemiology, Universitätsklinikum Tübingen, Tübingen, Germany

**Keywords:** GWAS, Python, summary statistics, visualization, quality control

## Abstract

There has been an exponential increase in the development of technologies that have offered unprecedented opportunities to unravel the genetic underpinnings of rare and complex diseases. However, despite the progress, and availability of tools, there are still several challenges to overcome, such as the development of a unified framework that facilitates downstream analysis. To address this challenge, we present IDEAL-GENOM (Integrated Downstream Analytical Toolkit for Genomic Analysis), a Python-based wrapper designed to streamline the analytical workflow commonly implemented in genome-wide association studies (GWAS) settings. IDEAL-GENOM integrates widely used tools such as PLINK, GCTA and bcftools along with custom-developed functionalities, enabling reproducible results through parameter sharing. IDEAL-GENOM provides a simplified framework for quality control (QC) and GWAS and downstream analysis that will enable beginners and advanced users to leverage the in-built functionalities to perform GWAS analysis for complex diseases. IDEAL-GENOM is a customizable tool that enables users to efficiently analyze genotyping data with reproducible QC, post-imputation, and GWAS pipelines.

## 1 Introduction

The advancement of genomic technologies along with simultaneous growth in computational capabilities —including fast processors and expanded memory capacity— has enabled large-scale population-based genetic studies to explore the genetic architecture for complex diseases with unprecedented depth (Loos 2020).

Despite the progress, because of the processing of large samples, various factors such as a less-than-optimal level of data quality coupled with low coverage can impact the results [Anderson 2010, Turner 2011]. This has a serious impact on running genome-wide association studies (GWAS) analytical pipeline, which has become a bona fide approach to identify potential novel genetic risk loci for complex diseases (Visscher, Brown, et al. 2012; Visscher, Wray, et al. 2017; Abdellaoui et al. 2023).

Various computational tools have been developed to streamline data management pertaining to datasets used for GWAS analyses. However, integrating these tools into cohesive, reproducible workflows remains a persistent challenge. For instance, while many QC steps can be performed using PLINK1.9, others require PLINK2.0 or a mix of scripts written in programming languages such as Perl or R. Similarly, GWAS can be performed using either PLINK or GCTA, but down-stream analyses often necessitate combining multiple tools in an unstructured manner (Purcell, Neale, et al. 2007; Yang et al. 2011; C. C. Chang et al. 2015; Purcell and C. Chang 2024). This fragmented approach complicates reproducibility because manually combining scripts can lead to error propagation (Roth et al. 2025).

To address these challenges, we introduce IDEAL-GENOM, a free and open-source Python library designed to integrate and streamline genomic data analysis. The analytical workflow is enriched with multiple visualization functionalities to allow users to control how the pipeline behaves. Our toolkit minimizes dependencies in the sense that it is completely written in Python, while offering customization, publication-ready plots and easy-to-use example notebooks to guide users through its functionalities.

By emphasizing simplicity, flexibility, and reproducibility, IDEAL-GENOM empowers researchers to focus on generating insights rather than navigating technical hurdles. It also constitutes an easy-to-use tool for beginners as well as for advanced users. Even though the design is primarily intended for case/control studies in human populations, we think that it might be adapted to another species as well.

## 2 Methods

### 2.1 Standard Pipelines

IDEAL-GENOM QC pipeline was designed following the protocol presented in (Anderson et al. 2010). This pipeline consists of sample-level QC, ancestry check, and variant-level QC. At the sample-level, the filters considered are call rate, sex concordance, heterozygosity, and relatedness. For ancestry check, we have developed the pipeline for ethnicity homogeneous populations, that is, populations with no admixture and that could be compared to the 1000 Genomes Project (1KG) superpopulations. While at the marker level, the filtering criteria are for nonrandom missingness by haplotype, Hardy-Weinberg Equilibrium (HWE) deviations in controls, and variant-level genotype missingness. The input and output files are PLINK1 binary (bed, bim, fam) files, but we intend to expand the accepted files to include PLINK2 binaries as well. In addition to the standard case/control pipeline, there is a complementary step where the population structure is studied in more detail. The GWAS pipeline consists of two stages, the first one is dedicated to processing and harmonizing imputed genome data, and the second is properly dedicated to GWAS analysis.

To facilitate reproducibility of the entire QC process, all parameters are stored in a YAML configuration file. This file provides full control over the pipeline steps to be executed, as well as the PLINK parameters. We recommend consulting its structure on GitHub. Moreover, the default values are based on our experience and are not intended as universal recommendations for all study scenarios.

#### 2.1.1 Quality Control

The pipeline is designed to minimize human intervention, requiring only a few command line interface (CLI) parameters to be executed. The hierarchically organized pipeline operates on a per-project basis, meaning that each cohort requires its own dedicated directory to ensure proper functionality (more details can be found in the documentation), and enhance data traceability. By adhering to this structure, the pipeline not only facilitates smoother data management but also supports reproducibility, ensuring that the entire analysis process can be accurately reconstructed and verified.

The QC pipeline consists of three steps: sample QC, ancestry check, and variant QC, which can be complemented with a subroutine intended to give a more detailed vision of the population structure, that is to apply two non-linear dimensionality reduction techniques: Uniform Manifold Approximation and Projection (UMAP) and *t*-distributed Stochastic Neighbor Embedding (*t*-SNE) to grasp the non-linear structure of the genome, and the *F*_*st*_ statistic computation of the cohort under study with the 1KG superpopulations (Figure 1A) (Healy et al. 2024; Maaten et al. 2008). The output of the pipeline is a cleaned PLINK binary file that could be used for different analysis, such as a deeper population structure analysis, a GWAS, or an imputation pipeline. In addition to the cleaned files, a variety of plots are generated as reports of the execution steps, such that the user may change the default parameters according to its needs.

**Figure 1:**
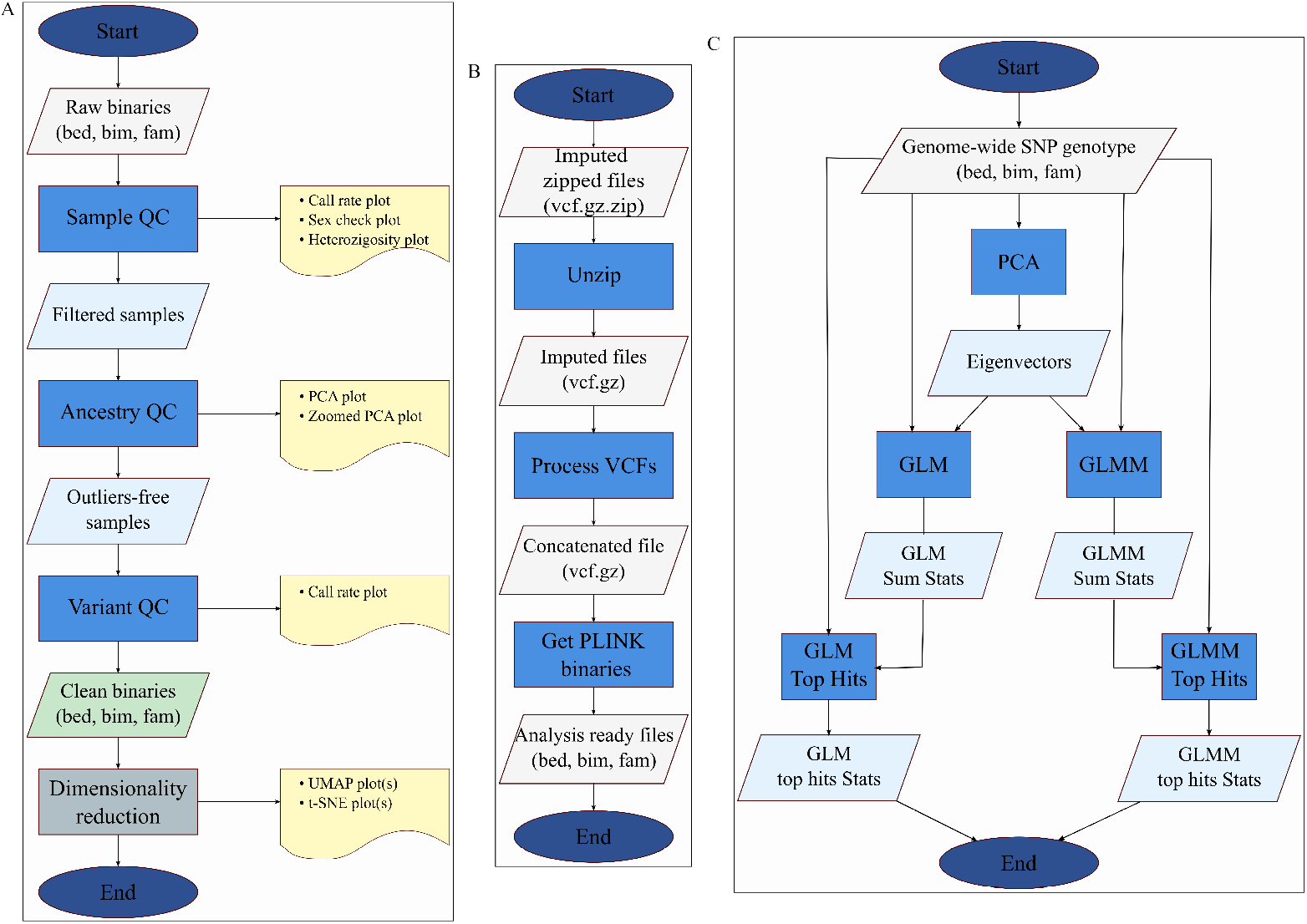
**A**. Quality control workflow **B**. Post-imputation workflow **C**. GWAS workflow.

#### 2.1.2 GWAS Pipeline

IDEAL-GENOM’s GWAS pipeline was designed as a “forked” pipe, so it can be viewed as a standard wye pipe. We have decomposed the workflow into four steps, with the goal of a more modular and versatile functioning that could also be integrated into different processes. The steps of the pipeline are post-imputation processing, pruning and PCA decomposition, association analysis in terms of a generalized linear model (GLM) and association analysis in terms of a generalized linear mixed model (GLMM). Once the summary statistics are ready, a stepwise model selection of independently associated SNPs is run with GCTA to obtain top hits among the significative SNPs. In addition, we have included an additional step to automatically annotate top SNPs with the nearest gene name (Figure 1C).

We also designed a subroutine that could be incorporated before the GWAS pipeline; that is the processing of imputed genomic data as password-protected zip files. It is common to have a cohort of genotyped data, and after QC submit the cleaned data to an imputation server. Usually, the output is a set of zipped vcf files that require preprocessing to perform GWAS. Hence, this subroutine can be incorporated as a step prior to GWAS, where the output is also a set of PLINK binaries (Figure 1B). This subroutine is implemented using PLINK1.9, PLINK2.0 and bcftools as building blocks (Danecek et al. 2021).

#### 2.1.3 2.1.3 Ancestry Check

We use the 1000G as our reference panel for the ancestry check. Our pipeline includes functionality to automatically download the reference panel (GRCh38 by default) and process it to generate the binary files required for ancestry analysis. The built-in functionality of our pipeline automatically identifies duplicate rsIDs within the 1000G dataset and standardizes SNP identifiers by renaming them to the format `chr:pos:a2:a1.` Thus, our tool automatically mitigates the exact matching of SNP identification and saves considerable user time in resolving the issue of SNP matching for ancestry check.

Our approach to detecting population outliers is simple and conservative: (1) Data alignment: the study dataset is first aligned with the reference panel through preparatory steps; (2) Principal Component Analysis (PCA): The pipeline computes the principal component decomposition of the dataset resulting from the merging of the study data and reference panel, which is complemented with a scree plot and cumulative explained variance for diagnosis; (3) Outlier detection: a) Samples falling outside a neighborhood (radius equal to `ref _threshold` times standard deviation and Chebyshev metric) of the centroid of the reference panel superpopulation are flagged as outliers, b) Samples falling outside a neighborhood (radius equal to **pop threshold** times standard deviation and Chebyshev metric) of the centroid of the cohort are flagged as outliers; (4) the final set of population outliers consists of the intersection of both groups.

Even though the sample size can affect the outlier detection method b), because the mean and standard deviations are less robust estimators, the method a) is more robust because it relies on the reference panel. Since we consider the intersection of methods, the methodology is robust to variations in sample size. Moreover, the user can change the metric to determine the centroid neighborhood and use another Minkowski metric instead of Chebyshev.

#### 2.1.4 2.1.4 Visualizations

The QC pipeline is complemented with a comprehensive set of performance report graphics designed to help the user diagnose each step. These allow the user to identify appropriate thresholds and parameters. The dimensionality reduction subroutine generates 2D plots for each combination of parameters explored, enabling the user to visually assess population structure and select the most informative configuration. Since UMAP and *t*-SNE are stochastic by nature, reproducibility is ensured by allowing the user to set a random seed, guaranteeing identical results across independent runs. There is also an extensive set of functions to visualize GWAS summary statistics, such as Manhattan, Miami, Trumpet, Q-Q, and effect-size comparison plots, each addressing a different aspect of the results interpretation.

## 3 Results

### 3.1 QC Pipeline Benchmark

IDEAL-GENOM is the genotype processing pipeline for the LuxGiant consortium (Kishore et al. 2025), for benchmarking it was used 1KG dataset. A Lenovo ThinkStation P3 Ultra on Ubuntu 24.04 3 LTS operating system and Intel Core i9-14900 x32 with 64 Gb of RAM was used. We compared our results with two existing tools: GenoTools and plinkQC but since the ancestry check subroutines are different in the three cases, we compared only the equivalent steps: sample and variant level QC (Vitale et al. 2025; Syed et al. 2025). The input was PLINK bed, bim, fam binary files as well as the output.

We ran the pipeline ten times for each software, and the results can be seen in Table 1. GenoTools average elapsed time was 57:30 minutes, while plinkQC average was 4:15:58 hours and IDEAL-GENOM averaged 53:00 minutes. These results show an improvement of 7.81% respect to GenoTools and 79.29% respect to plinkQC. Moreover, GenoTools and IDEAL-GENOM make better use of PLINK parallel computing possibilities.

**Table 1.**
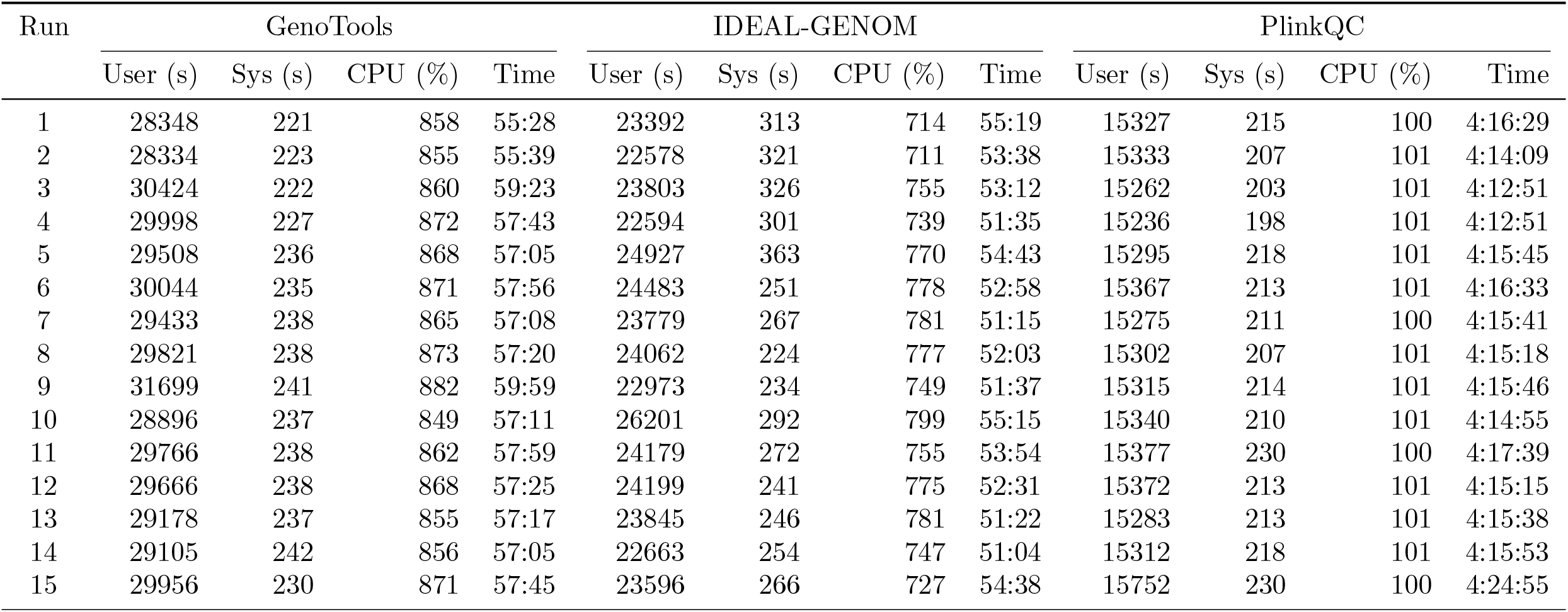
Performance comparison of GenoTools, IDEAL-GENOM, and PlinkQC across runs.

IDEAL-GENOM has helped to the discovery of new loci associated to Parkinson’s disease in Indian population, streamlining QC and GWAS pipeline as well as generating publication ready visualizations, reducing researchers’ burden of dealing with technical issues.

### 3.2 GWAS Pipeline Benchmark

To the best of our knowledge, there is no similar tool to IDEAL-GENOM for the GWAS pipeline. For example, plinkQC only performs quality control, while GenoTools only incorporates PLINK’s GLM model into its pipeline. In addition, GenoTools relies solely on kinship estimation for relatedness/duplicate check, and plinkQC depends only on identity-by-descend estimation, while IDEAL-GENOM allows to select which method is better suited for the available data. Consequently, IDEAL-GENOM offers a more user-oriented, flexible framework for GWAS analysis. Of note, when comparing GWASlab, a Python library designed to create high-quality plots from GWAS summary statistics, our tool offers a more modular and robust implementation He et al. 2023.

A subroutine is planned for the next release to automate imputation pipelines as well as adding functionalities to assess population structure and perform polygenic risk score computation.

## 4 Discussion

IDEAL-GENOM toolkit has been developed to offer a user-friendly framework to provide training to beginners and advanced users for the Lux-GIANT consortium Rajan et al. 2020; Kishore et al. 2025. The toolkit enables researchers to conduct end-to-end genomic analyses, from quality control to GWAS visualization, within a single reproducible framework. This Python wrapper has encapsulated QC and ancestry check workflows with post-imputation and pre-GWAS data wrangling. Consequently, many time-consuming steps have been unified in a seamless workflow with multiple customization parameters. It allows researchers to obtain robust and trustful results, where these analyses can be easily reproduced. IDEAL-GENOM continues to be developed and maintained and intends to become an open-source and collaborative project.

## Data Availability

Data from 1000G project is freely available at https://www.internationalgenome.org/data and https://www.cog-genomics.org/plink/2.0/resources#phase3_1kg. All the code is available on GitHub https://github.com/cge-tubingens/IDEAL-GENOM-QC. The library is also available on PyPI https://pypi.org/project/ideal-genom/ and its documentation is on Readthedocs https://verus-ideal-genom.readthedocs.io/en/latest/.

